# Choosing optimal trigger points for *ex situ, in toto* conservation of single population threatened species

**DOI:** 10.1101/2021.01.08.425867

**Authors:** Kaitlyn Brown, Tamara Tambyah, Jack Fenwick, Aidan Burke, Patrick Grant, Solene Hegarty-Cremer, Joshua Muller, Michael Bode

## Abstract

Many endangered species exist in only a single population, and almost all species that go extinct will do so from their last remaining population. Understanding how to best conserve these single population threatened species (SPTS) is therefore a distinct and important task for threatened species conservation science. As a last resort, managers of SPTS may consider taking the entire population into captivity – *ex situ, in toto* conservation. In the past, this choice has been taken to the great benefit of the SPTS, but it has also lead to catastrophe. Here, we develop a decision-support tool for planning when to trigger this difficult action. Our method considers the uncertain and ongoing decline of the SPTS, the possibility that drastic *ex situ* action will fail, and the opportunities offered by delaying the decision. Specifically, these benefits are additional time for ongoing *in situ* actions to succeed, and opportunities for the managers to learn about the system. To illustrate its utility, we apply the decision tool to four retrospective case-studies of declining SPTS. As well as offering support to this particular decision, our tool illustrates why trigger points for difficult conservation decisions should be formulated in advance, but must also be adaptive. A trigger-point for the *ex situ, in toto* conservation of a SPTS, for example, will not take the form of a simple threshold abundance.

## INTRODUCTION

The Earth’s most threatened species are often only found in a single location. Some were originally range-restricted endemics, while others have suffered catastrophic range and abundance declines. Well-known examples of “single population threatened species” (hereafter, SPTS) include the last remaining Javan rhinoceroses *Rhinoceros sondaicus*, which are restricted to a single Indonesian national park, Ujung Kulon [1]. Similarly, the Annobón scops owl *Otus feae* is found only on the island of Annobón in Equatorial Guinea [2], and the distribution of the Hainan gibbon *Nomascus hainanus* has shrunk to a single nature reserve in China [3]. The Alabama cavefish *Speoplatyrhinus poulsoni* only exists in a single cave in the Key Cave National Wildlife Refuge [4], while the global distribution of the Fitzroy Falls crayfish *Euastacus dharawalus* in Australia has dwindled to less than a square kilometer of habitat [5].

SPTS face a particularly high risk of extinction because even local threats can cause global extinctions. It is a central and foundational tenet of conservation biology that species with multiple populations have much greater insurance against global extinctions [6,7]. Remaining in a single population is so threatening – and so common – that the IUCN Red List automatically categorises SPTS as Critically Endangered, its highest category of risk. Natural population fluctuations can push populations to low abundance, where they become vulnerable to Allee effects, genetic bottlenecks, or simple misfortune. Even SPTS with high abundance and genetic diversity can still be threatened by environmental catastrophes. The Javan rhino, for example, could be pushed to extinction by tsunamis generated in nearby Anak Krakatoa [8]. SPTS are particularly vulnerable to human activities, even when protected by legislation. Australia’s stocky galaxias *Galaxias tantangara* (a listed species) are only found in the headwaters of Tantangara creek, and the approved development of hydroelectric infrastructure is likely to introduce competitively superior species from adjacent catchments [9]. Although not all SPTS are currently in decline, most will experience periods of decline at some stage. Without conservation interventions, some SPTS will become extinct.

Management actions for SPTS can be broadly classified as either *in situ* or *ex situ* [10]. *In situ* conservation actions attempt to improve population viability within a species’ natural environment. *Ex situ* actions remove the species from this habitat (and therefore from the threatening process), generally into a captive breeding program. While both strategies may be undertaken simultaneously, for SPTS this choice is often exclusive. Moreover, when declines are rapid and abundances are low, it may be necessary to take the entirety of the remaining population into captivity. This extreme form of *ex situ* management – *ex situ, in toto* – is controversial because the species immediately becomes extinct in the wild. For this reason, it is only implemented as a last resort [11].

*Ex situ, in toto* conservation decisions are a genuine conservation dilemma. The action often appears essential, given a small, shrinking population. However, it is drastic step, with a mixed record of success. In some cases, *ex situ, in toto* actions were able to pull SPTS back from the precipice of extinction. The California condor *Gymnogyps californianus* is perhaps the most famous example: in 1987, after decades of dwindling numbers, the remaining population of 27 individuals were taken into a captive breeding program. Their abundance recovered and the species was successfully re-released into the wild, where it now numbers 312 individuals (with 176 additional condors in captivity; USFWS, 2018). Other examples of successful *ex situ, in toto* actions followed by successful reintroduction include red wolves *Canis rufus* (also *C. lycaon rufus* or *C. lupus rufus*, depending on the authority); the Guam kingfisher *Todiramphus cinnamominus* and Guam Rail *Hypotaenidia owstoni*; and black-footed ferrets *Mustela nigripes*.

For other species, an *ex situ, in toto* actions hastened or caused their extinction. In South Australia in 1924, government scientists attempted to take the last known individuals of the toolache wallaby *Macropus greyi* into captivity. During the translocation, 10 of the 14 remaining individuals were accidentally killed and the remaining 4 re-released. The depleted population went extinct by 1939. For a final set of SPTS, the opportunity to undertake *ex situ, in toto* action passed before managers could make the difficult decision. This situation – monitoring a species to extinction [13] – is depressingly common in conservation. The Christmas Island Pipistrelle *Pipistrellus murrayi*, for example, was common until 1984, but thereafter began to decline for unknown reasons. Captive breeding was proposed as early as 2006, but was delayed for additional research until July 2009. By this point, no individuals could be found in the wild.

Retrospective analyses of the Pipistrelle extinction [14], and that of other threatened species extinctions [13,15], suggest that many conservation failures – including some of the examples above – are partly attributable to inadequate preparatory planning. In particular, these analyses recommended that managers need to anticipate extinctions, and should identify thresholds (“trigger points”) for when important actions should be implemented. Trigger points offer two significant benefits. First, they proactively use the best information to calculate when to act, weighing the relevant costs, benefits, and uncertainties [15]. Second, they insure decision-makers against psychological factors such as political pressure or shifting-baselines [16], which might cause managers to shy-away from a difficult decision at the critical moment.

While there are urgent calls for managers to set explicit trigger-points for drastic action when threatened species are in decline [13,14], there is no theoretical guidance available to support this decision for SPTS. The decision to initiate *ex situ, in toto* actions combines high-stakes and high-uncertainty. It is critical that managers make the correct choice, but the irreversible and uncertain nature of the action makes it a very difficult decision. Our goal in this paper is to design a simple and robust decision-support tool that can help managers set a trigger point for *ex situ, in toto* action for a SPTS.

## METHODS

Trigger points for *ex situ, in toto* conservation of SPTS should incorporate three essential decision factors. The first and overriding concern is with uncertainty. Conservation decisions always involve uncertainty, and this is particularly true for the choice to initiate *ex situ, in toto* conservation. Uncertainty enters the decision at multiple points. There is uncertainty about the rate of population decline, which can be difficult to measure accurately. There is also uncertainty about whether ongoing *in situ* conservation actions will be able to halt the decline [17]. Finally, there is uncertainty about whether a captive breeding program – if attempted – will successfully establish an *ex situ* population [18–20].

The second complexity is the time-dependent nature of the decision. While a decision to delay action risks further population decline, it also gives ongoing *in situ* actions another opportunity to succeed. Decisive *ex situ* actions reduce future flexibility, and this cost must be incorporated in the decision. The third complexity involves learning (i.e., the reduction of uncertainty): until *ex situ, in toto* actions are triggered, managers can observe and learn about both the speed of population decline, and also the probability that *in situ* actions will be successful. Both pieces of information mean that longer periods of *in situ* management will be associated with better decisions. Setting trigger points is thus also a choice about how much learning will be allowed [17,21,22].

We therefore formulate the trigger point decision as a Bayesian optimal stopping problem, based on Markov chain model of the system dynamics [23,24]. This formulation allows us to incorporate uncertainty, dynamical decisions, and learning, and to solve for the optimal solution using stochastic dynamic programming (SDP; Bellman, 1954; Bode & Possingham, 2007).

We assume that the abundance *N_t_* of a SPTS has been observed for *t*_0_ years, during which time it appears to have declined at a linear rate (Figure 1 shows a hypothetical example; Figure 2 shows historical case-studies). Fitting this data with a regression line yields a probability distribution *p*(*r*) for the decline rate, which allows us to construct a probability distribution for the expected time to extinction *p*(*t_E_*) in the absence of successful *in situ* or *ex situ, in toto* actions. While this distribution may not be bounded (e.g., if we assume that the observation error is Gaussian), in practice we will choose an upper limit *τ* (e.g., the upper 99^th^ percentile), beyond which we assume extinction is effectively guaranteed (a finite time horizon is needed to implement SDP). Note that we are assuming that variation in its observed decline trajectory is due to environmental and demographic stochasticity, not sampling error.

**Figure 1:**
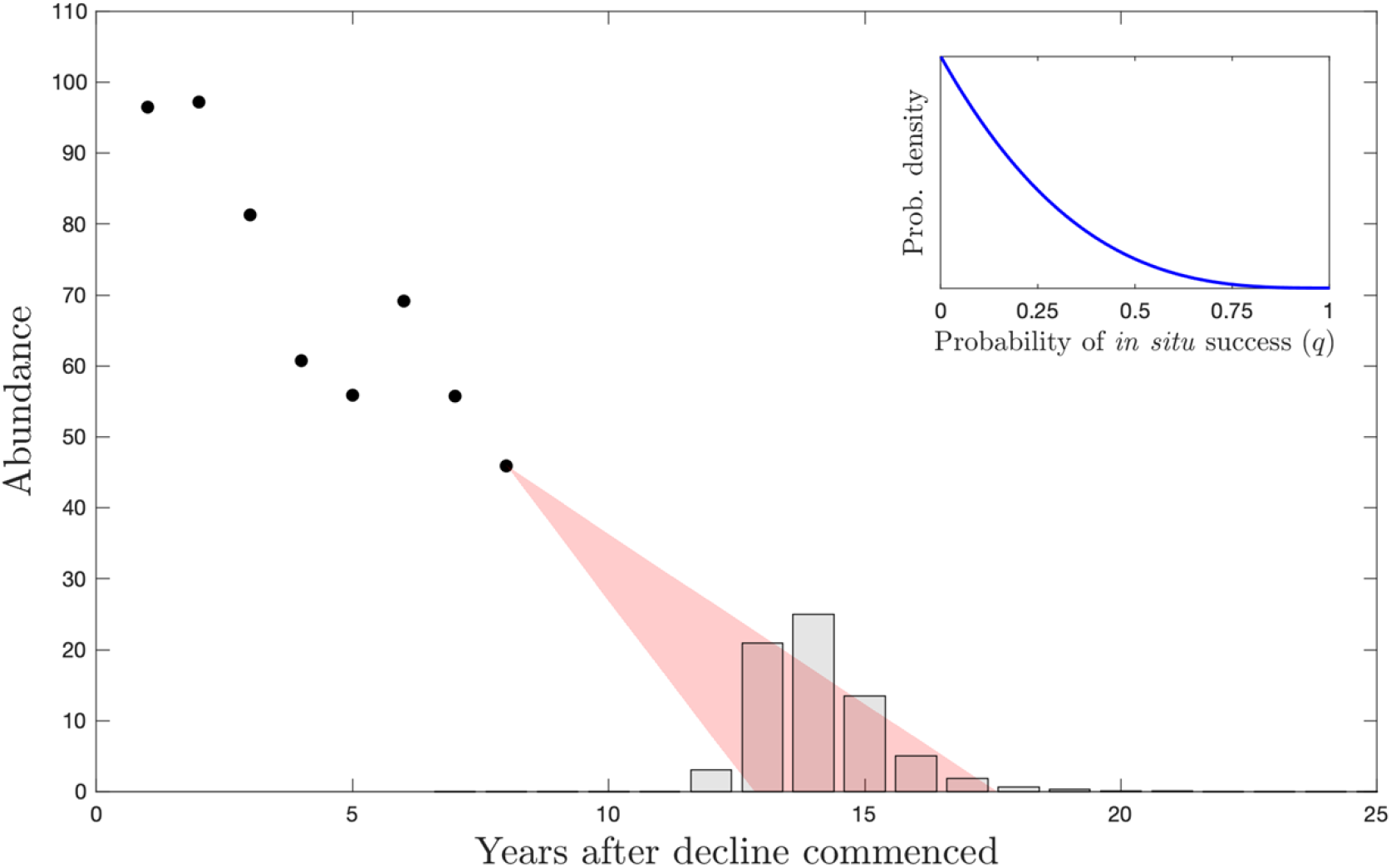
Schematic of an *ex situ, in toto* decision for a hypothetical single population threatened species (SPTS). The SPTS abundance has been observed to decline at a linear rate for 8 years; extrapolating, the population will likely drop to extinction within 4 – 10 years. The bars show the probability distribution for the extinction year, based on the range of linear declines shown in red. Conservation actions have been underway for 3 years without success, yielding a belief distribution in the annual probability of success that is shown in the inset plot.

**Figure 2:**
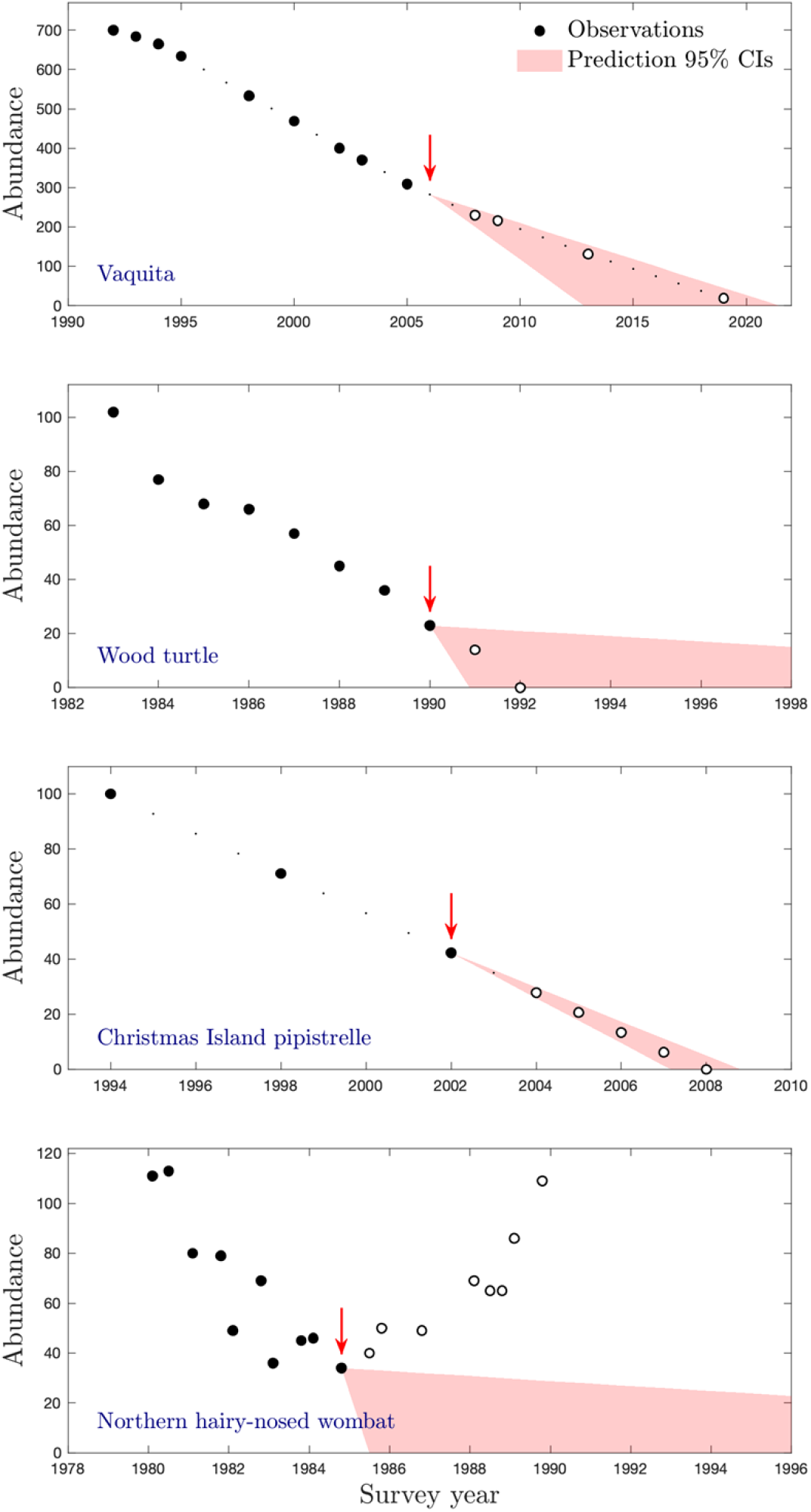
Results of applying the optimal decision method to four declining case-study populations. In each case, the red arrow shows the point in time at which the method recommended *ex situ, in toto* action be undertaken, based on the range of linear declines shown in red. Closed markers show observations preceding the decision, open circles show observations after the decision would have been recommended.

At some point in the past, conservation managers began to apply *in situ* conservation actions. These might be a single consistent action, a sequence of different actions, or a suite of actions all at once. We assume that these *in situ* actions will successfully reverse or halt the observed ongoing decline with an unknown (but constant) probability *q.* The probability *q* is not known with certainty, and may be zero. Our estimate of this parameter will improve through time, as managers observe the failure or success of the action. Repeated failures will make the managers increasingly pessimistic about the value of *q*, and therefore the likelihood that the action will eventually work (Figure 1 inset). In particular, if we assume that the SPTS response to the conservation action is a Bernoulli trial, then standard results from adaptive management [27] allow us to calculate the posterior belief about the value of *q.* If *in situ* management actions have been unsuccessful for *n_u_* years, then the probability that the next year’s attempt will be successful is the beta distribution *f*(*x*;*α* = 1, *β* = *n_u_* + 1). The expected value of *q* is 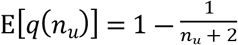.

Managers also have the option of applying a single *ex situ, in toto* action, which has probability of success *p_x_*. This parameter is clearly critical to the decision, but there will rarely be data available to estimate its value. Moreover, since we are considering a small SPTS, we assume that there is no opportunity to learn about *p_x_* over time. Canessa and colleagues [28] suggest that expert elicitation be used to estimate its value. Alternatively, Dolman and colleagues [29] used a stochastic, individual-based population model.

We can formulate the dynamics of this decision problem as a discrete time Markov chain model, with system states defined by three factors. First, the number of years *n_u_* that the *in situ* action has been attempted unsuccessfully (a number between 0 and *τ*); second, whether an *ex situ, in toto* action has been taken (a binary variable *α_e_*); third, whether the *in situ* action has been successful (a binary variable *a_s_*). Thus the system is initially in state {*n_u_,a_e_,a_s_*} = {0, 0, 0}, if the managers have not previously attempted the *in situ* management action, and therefore have no information about its probability of success. If the state were {3, 0, 1}, then four years have elapsed since *in situ* management began, *ex situ, in toto* actions were not taken, and in the fourth year the *in situ* management succeeded in reversing the population decline. Note that, because we assume that *in situ* and *ex situ, in toto* actions cannot both be undertaken in the same year, the number of years that have elapsed in the project is the sum of the values in the state vector. We uniquely enumerate state *S_i_* = {*n_u_,a_e_,a_s_*} with the index *i* = *a_s_* +2*a_e_* +4*n_u_* +1, and there are a total of *R* = 4(*τ* + 1) states.

The system dynamics are expressed by two probabilistic transition matrices: **T**^(*s*)^ and **T**^(*e*)^. The former matrix describes the state dynamics that occur when the managers undertake *in situ* actions, and the latter describes the dynamics when managers decide to implement the drastic *ex situ, in toto* action. We can define the elements of the *in situ* matrix as:

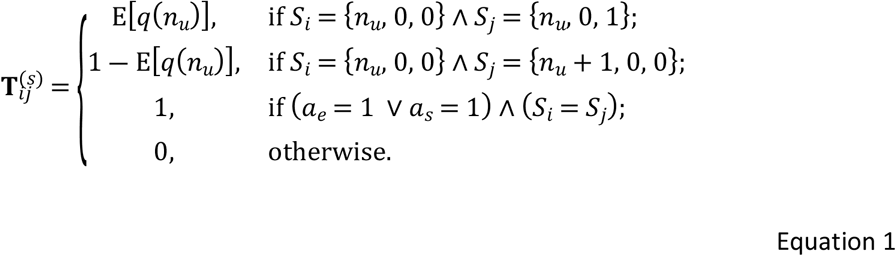

The first case describes the state dynamics associated with a successful *in situ* action, the second with an unsuccessful *in situ* attempt. Note that these transition probabilities are state-dependent (specifically, on *n_u_* the number of years that *in situ* management has been unsuccessful), and that managers are therefore able to anticipate future learning. The third case reflects the fact that once either *ex situ, in toto* action has been taken, or an *in situ* action was successful, the system no longer undergoes transitions (i.e., it has reached an absorbing state). Finally, we assert for completeness that the transition from state *S_i_* = {*τ*, 0, 0} to itself occurs with probability 1.

We can define the elements of the *ex situ, in toto* matrix as:

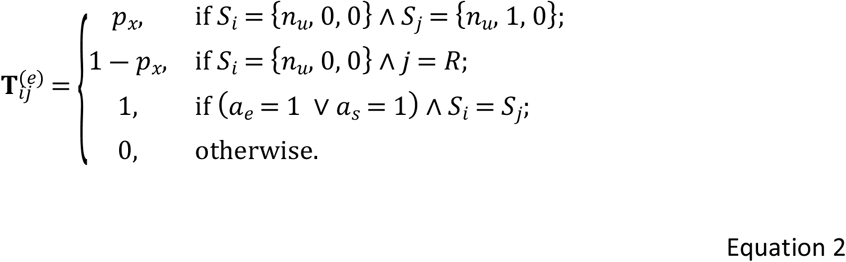

In this matrix, the first case describes the state dynamics associated with a successful *ex situ, in toto* action. In the second case, an unsuccessful *ex situ, in toto* intervention resulted in the species becoming entirely extinct, which we model as a transition to the final state of the model *S_R_* = {*τ*, 1, 1}. This is a convenience that we can apply without altering the state dynamics, since this state cannot be reached from the initial state. The third case once again reflects the two categories of absorbing state.

It can be helpful to calculate the expected abundance of the SPTS in each of the system states. While this value is not included explicitly in the system state, it can be estimated on the basis of the observed (uncertain) population decline and the final observed abundance *N*(*t*_0_):

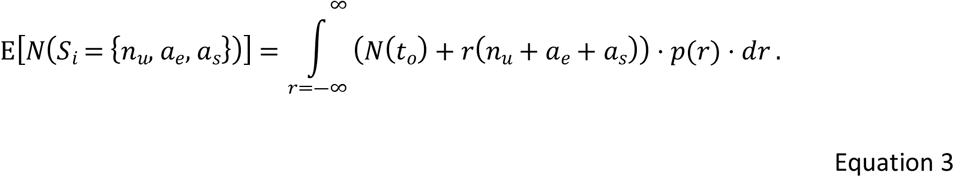

This equation relies on the fact that the total time elapsed is the sum of the number of unsuccessful attempts at *in situ* actions, and the success state of *in situ* and *ex situ, in toto* actions. Because the successful states are absorbing (see Equations 2–3), the summation (*n_u_* + *a_e_* + *a_s_*) captures the time until success.

Our conservation objective for a SPTS is to ensure its persistence. To apply SDP, we first need to define the relative value of being in each system state at the terminal time τ. By this time, the final state of the SPTS will be definitely known: either persisting *in situ*; persisting *ex situ* and extinct in the wild; or completely extinct. This value function should be defined for each particular threatened species. Here, we construct a general version of this function that reflects two factors. First, the value should be positively related to the abundance of the SPTS at the time that the *in situ*, or the *ex situ, in toto* action succeeds. Smaller population sizes are more exposed to extinction, and have less genetic variation, even as they begin to recover. Larger population sizes are worth more, but increasing abundance should deliver diminishing marginal returns. Second, the value of a state should depend on whether the SPTS is *in situ*, or *ex situ, in toto* (with the former being preferable). We therefore define the value function as:

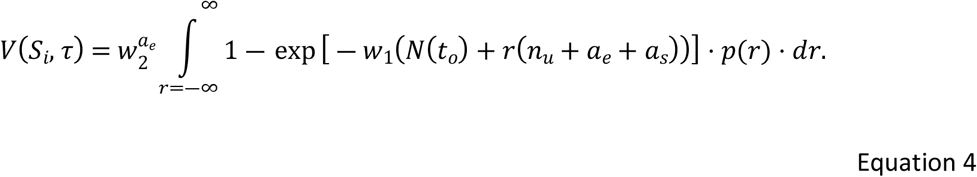

In this equation, *w*_1_ controls the exponentially diminishing marginal returns of larger populations, diminished by a factor *w*_2_ if that population is secured *ex situ, in toto*, rather than *in situ.*

Armed with the transition matrices and value function, stochastic dynamic programming can be applied via backward iteration, to determine the optimal action to take, in every state and year. The expected value for each system state, at intermediate times, can calculated using the stochastic Bellman equation:

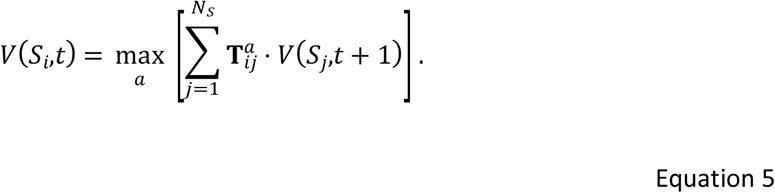

Following Bellman’s principle of optimality, we identify the best management policy by repeatedly iterating this equation, and recording each state- and time-dependent optimal action *a*\*(*S_i_,t*).

### Case studies

To illustrate the application of the method and its results, we use it to assess 4 historical datasets of declining vertebrate populations: the vaquita *Phocoena sinus*, the Christmas Island pipistrelle, the northern hairy-nosed wombat *Lasorhinus krefftii*, and the North American wood turtle *Clemmys insculpta* (Figure 2). We choose these datasets because they are published examples of declining populations with well-studied abundance timeseries, and because they exhibit a range of final outcomes, including extinction. The vaquita population has shown more than 25 years of ongoing decline, and currently numbers fewer than 50 individuals. The pipistrelle and turtle populations both declined monotonically to extinction. Finally, the wombat population recovered after a period of precipitous decline, after *in situ* management actions and environmental changes saved the species. See *Supporting information* for more details on the data, and optimal solution to each of these case study populations.

For each case-study, we choose parameters for the value function that are arbitrary but plausible (for real SPTS, we expect that these parameters would be chosen by ecologists and stakeholders). First, we assume that managers’ preference for wild *in situ* populations over captive *ex situ, in toto* populations is *w*_2_ = 2/3. Second, we assume that managers have a relatively high satisfaction with the initial conditions of the *in situ* population, before the onset of decline (specifically, that they were 95% satisfied, and therefore that *w*_1_ = ln(20)/*N*(*t_o_*)). Finally, we assume that *ex situ, in toto* action is more likely than not to be successful (*q* = 0.75), but that there is still a substantial probability of failure.

We apply our SDP optimal solution method to each dataset in turn, simulating the actions of a decision-maker who begins to take *in situ* actions at in the first year of the population decline. Our simulated optimal decision processes assume that the dataset is being gathered in real time. After each new abundance data point is gathered, we re-estimate the decline rate, update the posterior probability of *in situ* success, and calculate whether the decision-maker should initiate *ex situ, in toto* conservation. We continue this process until either the SDP solution recommends that *ex situ, in toto* action should be taken, or extinction occurs.

## RESULTS

The timeseries for each of the case-study populations is shown in Figure 2, and the SDP-defined solution to the *ex situ, in toto* conservation problem has been superimposed on these observations.

The circular markers denote the observed timeseries. The red arrows denote the point at which the SDP solution recommended that *ex situ, in toto* conservation action be taken The red shaded region shows the 95% confidence intervals of the predicted future SPTS abundance at that point. Uncertainty varied between the different case-studies, with the wood turtle and wombat timeseries exhibiting the greatest abundance variation, and the vaquita and pipistrelle timeseries both showing consistent and linear declines over more than a decade.

In all four cases, the optimal solution triggers *ex situ, in toto* conservation action before the population declines to extinction. In the case of the Christmas Island pipistrelle, *ex situ, in toto* actions are recommended 6 years before the population declines to zero. For the vaquita, the optimal solution recommends that dramatic *ex situ* actions should be taken in 2006, more than 15 years ago. *Ex situ, in toto* actions are recommended for the wood turtle, but they only in 1990, a mere two years before the population vanishes. Finally, for the northern hairy-nosed wombat, the optimal solution recommends *ex situ, in toto* conservation of the SPTS in 1985, the precise year that the population began to recover.

## DISCUSSION

As we outlined in our introduction, extinction is an ever-present danger for any of the Earth’s many SPTS. Experience teaches us that conservation can often fail to intervene effectively when SPTS have declined in the past, and that a key driver of this inaction is a lack of effective planning. We have therefore formulated a straightforward, optimal stochastic management tool for a declining SPTS.

In our optimal solutions (Figure 2), the decision about whether to take or delay *ex situ, in toto* action balanced several competing factors. The superiority of *in situ* populations over *ex situ* is a constant factor recommending against intervention. However, the relative belief in the probability of *in situ* success (*q*, which declines through time) versus *ex situ* success (*p_x_*, which remains constant) diminishes through time, until *ex situ* actions deliver superior expected outcomes. This process of learning, which will occur in all ongoing conservation management projects, means that trigger points must be more complicated than simple abundance thresholds (e.g., *“Ex situ* action should be taken one the population falls below 100 individuals”). In Figure 2a, for example, the optimal manager would have taken action once the population fell below 300 individuals. However, this trigger point is specific to a timeseries which began (along with unsuccessful *in situ* management) in 1991. If observations and actions had only been started in 2000, by contrast, the trigger point for *ex situ, in toto* actions would have been lower.

Our approach can be readily applied to any declining SPTS, to ensure that decision-makers have an *a priori* plan for when to initiate the drastic step of *ex situ, in toto* conservation actions. In its current incarnation, the tool is not based on a sophisticated population model, and we would be happy to see it superseded by a more complex description of decline (e.g., an exponential function), or better still, a species-specific population viability model. For these reasons, our optimal solutions (Figure 2) are not intended as recommendations for these particular species, nor to retrospectively second-guess the conservation actions that were (or were not) taken. While we believe that our methodology can be productively applied to SPTS with this type of abundance data, several of our parameter assumptions and prior belief distributions could be improved with more information about the threatened species. In particular, the two w parameters in the value function will depend on how key stakeholders value *in situ* versus *ex situ* populations of different sizes. Similarly, our model of ongoing *in situ* actions, and their expected probability of success, could be superseded by a bespoke, parameterised model or by expert opinion [28,29]. The probability of *ex situ* actions being successful could be carefully estimated for the particular species, based on experience with similar types of species (e.g., several species in the genus *pipistrellus* have been successfully bred in captivity). Likewise, the probability of *in situ* actions succeeding could be given an informative prior, based on other information. Finally, we modelled the likely future abundance of the SPTS as a simple extrapolation of the observed decline. More complicated models could yield better predictions (e.g., by considering measurement error as well as ecological stochasticity), but these can be readily incorporated into the optimisation framework through different formulations of equations 3 and 4.

Despite these opportunities for improvement, for species that do not have such complex models, or for species where these models are still being constructed and parameterised, we believe that our tool offers a transparent and quantitative justification for taking (or not taking) a difficult and contentious conservation management action. Essentially, we believe that conservation management plans for every SPTS should contain a clear and defensible trigger-point for *ex situ, in toto* action, and we devised this tool to offer quantitative support for that decision.

